# Semi-Supervised Learning to Boost Cardiotoxicity Prediction by Mining a Large Unlabeled Small Molecule Dataset

**DOI:** 10.1101/2024.05.25.595894

**Authors:** Issar Arab, Kris Laukens, Wout Bittremieux

**Affiliations:** Department of Computer Science, University of Antwerp, 2020 Antwerp, Belgium; Biomedical Informatics Network Antwerpen (biomina), 2020 Antwerp, Belgium

## Abstract

Predicting drug toxicity is a critical aspect of ensuring patient safety during the drug design process. Although conventional machine learning techniques have shown some success in this field, the scarcity of annotated toxicity data poses a significant challenge in enhancing models’ performance. In this study, we explore the potential of leveraging large unlabeled datasets using semi-supervised learning to improve predictive performance for cardiotoxicity across three targets: the voltage-gated potassium channel (hERG), the voltage-gated calcium channel (Cav1.2), and the voltage-gated sodium channel (Nav1.5). We extensively mined the ChEMBL database, comprising approximately 2 million small molecules, then employed semi-supervised learning to construct robust classification models for this purpose. We achieved a performance boost on highly diverse (i.e. structurally dissimilar) test datasets across all three targets. Using our built models, we screened the whole ChEMBL database and a large set of FDA-approved drugs, identifying several compounds with potential cardiac channel activity. To ensure broad accessibility and usability for both technical and non-technical users, we developed a cross-platform graphical user interface that allows users to make predictions and gain insights into the cardiotoxicity of drugs and other small molecules. The software is made available as open source under the permissive MIT license at https://github.com/issararab/CToxPred2.

## 1. Introduction

Computational drug discovery is an interdisciplinary field at the intersection of computer science, chemistry, biology, and pharmacology, dedicated to accelerating and optimizing the process of identifying potential therapeutic compounds for treating diseases. Traditionally, researchers in drug discovery employ *in vitro* and *in vivo* studies to evaluate the pharmacodynamics and pharmacokinetic (PD/PK) properties of candidate compounds identified through preliminary screening [1][2]. However, these experiments are resource-intensive in terms of both time and finances [3][4][5]. Moreover, the use of animal testing in early drug discovery stages often sparks ethical debates [6]. Leveraging advanced computational algorithms, machine learning (ML) techniques, and high-throughput screening methods, computational drug discovery seeks to streamline the discovery, design, and optimization of novel drug candidates. By simulating molecular interactions, predicting pharmacokinetic properties, and analyzing large datasets, this field aims to expedite the identification of promising compounds while reducing the time, cost, and ethical concerns associated with traditional drug development approaches.

Toxicity stands out among the five key pharmacokinetic properties—further including chemical absorption, distribution, metabolism, and excretion—requiring thorough validation before approving a novel drug candidate for clinical trials [7]. As outlined by the directives of the International Conference on Harmonization of Technical Requirements for the Registration of Pharmaceuticals for Human Use, preclinical evaluations necessitate scrutiny of cardiac ion channels inhibition and QT interval (i.e. time duration between the start of the Q wave and the end of the T wave on an electrocardiogram) prolongation induced by small compounds [8]. This phenomenon, termed drug-induced cardiotoxicity, involves the inhibition of pivotal cardiac ion channels, namely the voltage-gated potassium channel (hERG), voltage-gated calcium channel (Cav1.2), and voltage-gated sodium channel (Nav1.5).

Drug-induced cardiotoxicity has emerged as a leading cause of drug withdrawals observed during post-market surveillance [9]. Over the past four decades, approximately 10% to 14% of withdrawals were attributed to concerns regarding cardiovascular safety, impacting previously successful treatments such as rofecoxib, tegaserod, sibutramine, and rosiglitazone, thus making it the third most prevalent cause of adverse drug reactions [10][11]. The pharmaceutical industry is facing a growing concern over the projected increase in drug withdrawals [10][12]. This is particularly worrisome due to the substantial losses already experienced from cardiotoxicities detected at various stages of drug development, prompting the removal of multiple drugs from the market or the suspension of numerous drug discovery programs [13]. Among other examples of affected therapeutic drugs are astemizole, terfenadine, sertindole, grepafloxacin, vardenafil, cisapride, and ziprasidone, which were either withdrawn or severely restricted due to their adverse cardiac toxicity side effects [14][15]16].

In recent years, there has been significant interest and widespread adoption of ML models for predicting the cardiotoxicity of small molecules. Such computational methods for predicting cardiac ion channels’ inhibition offer substantial advantages, especially in the initial phases of drug discovery to effectively screen out molecules with a high likelihood of failing in clinical trials. Several previous reviews [14][17][18][19] have underscored this shift, also highlighting a notable observation in recent ML methods presented, favoring the use of random forest (RF), support vector machine, and deep neural network methods, primarily due to their demonstrated superior empirical performance. As stated in our previous research [20], there exists significant room for improvement in the field of cardiotoxicity prediction. Such improvement includes expanding computational methods beyond solely focusing on hERG, but to also tackle the other crucial cardiac ion channels: Nav1.5 and Cav1.2. Additionally, using the optimal predictive features is essential, alongside establishing a standardized evaluation and benchmarking framework. Moreover, training models on large and carefully curated labeled databases is crucial for robust model development. However, the challenge of accessing such extensive databases persists due to the high cost and resource demands associated with labeling large datasets. Unlabeled data, on the other hand, represents an unexplored gold mine in drug discovery, offering a treasure trove of potential insights waiting to be unlocked. Its untapped richness lies in its capacity to reveal hidden patterns within the chemical and biological space, which can be leveraged for novel discoveries and transformative advancements in pharmaceutical research. In this work, we used it to improve cardiotoxicity models’ performance.

To the best of our knowledge, all models employed in the field heavily rely on supervised learning, and semi-supervised learning (SSL) has so far not yet been studied in cardiotoxicity prediction. The most related research to date has been conducted by Chen *et al*. [21], who applied the SSL paradigm in a broader context of toxicity prediction using the Tox21 dataset [22]. This dataset comprises twelve endpoints, encompassing seven nuclear receptor signals and five stress response indicators. Notably, none of these endpoints directly relate to cardiotoxicity or any cardiac function. The paper utilized the mean teacher SSL algorithm, which employs a weak augmentation strategy by perturbing the labeled data and injecting Gaussian noise into the molecular feature representation. While this approach is commonly used in computer vision [23][24][25], we posit that modifying either the descriptors or fingerprints could lead to entirely different compounds, potentially unrealistic molecules, or inadvertently overlap with the test sets, thus contributing unintentionally to data leakage in the modeling phase. Therefore, we opted for the adoption of a pseudo-labeling algorithm instead.

This study builds on our previous work [20] by mining the whole database of small molecules in ChEMBL, comprising approximately 2 million compounds. With this approach, our goal was to construct cardiotoxicity classifiers that are robust, reliable, and possess superior generalization capabilities. To optimally leverage these large unlabeled data, submodular optimization was employed to strategically sample a subset of 100,000 unlabeled small molecules deemed most representative for each ion channel target. We employed pseudo-labeling (also called self-training) for SSL model building. To make the software user-friendly to the wider audience, both technical and non-technical, we integrated the optimal models in a graphical user interface (GUI). The GUI provides an interactive way to generate insights on small molecules under development by computing a list of eight key physicochemical properties using the quantitative estimation of drug-likeness (QED) [26][27] module in RDKit (http://www.rdkit.org), the 2D structure, the cardiotoxicity predictions, and confidence prediction measures as a quantification of the models’ uncertainty.

## 2. Methods

### 2.1 Compilation of the Cardiotoxicity Development Database

The dataset used in this study consists of two categories: a labeled set and an unlabeled set.

In the labeled set, compounds exhibiting inhibitory activity were collected from multiple public data repositories, including the ChEMBL bioactivity database [28][29][30], PubChem [31], BindingDB [32][33], hERGCentral [34], and US patent and literature-derived data [35][36][37][38]. Further information on the manual data curation process can be accessed in [20].

After the compilation of the final unique labeled set, compounds were classified based on their IC50 values, following standard criteria used by researchers in the field [19][39][40][41][42]. Compounds with IC50 values of 10 μM or below (pIC50 ≥ 5) were categorized as blockers (inhibitors), while compounds with IC50 values higher than 10 μM (pIC50 < 5) were categorized as non-blockers (inactive).

In adherence to data science best practices for model development, we extracted two external/independent test sets from each target dataset (hERG, Nav1.5, and Cav1.2). We used RDKit (default settings) to compute the Tanimoto similarity [43][44] between each pair of compounds in the datasets using 2048-bit extended connectivity fingerprints, also referred to as circular or Morgan fingerprints [45]. The first test set consisted of compounds with a structural similarity of no more than 70% (Tanimoto similarity ≤ 0.7) to the remaining development set, while the second test set comprised compounds with a structural similarity of no more than 60% (Tanimoto similarity ≤ 0.6) to the remaining development set. These external unique sets were denoted as hERG-70 & hERG-60 for hERG, Nav-70 & Nav-60 for Nav1.5, and Cav-70 & Cav-60 for Cav1.2. (Refer to Supplementary Table S0 for the class distribution of blockers vs. non-blockers in each set and for each target ion channel).

Regarding the unlabeled data, the entire database of molecular compounds was extracted from ChEMBL (October 2023). This set comprised 2.2 million compounds, which underwent the following cleaning process. Initially, the data was filtered by “type” to retain only entries explicitly identified as a “small molecule”. Subsequently, the chemical structures within the dataset underwent standardization using the Python packages RDKit and MolVS (https://github.com/mcs07/MolVS). The standardization procedure involved selecting the largest fragment, removal of explicit hydrogens, ionization, and stereochemistry calculation. The compounds were represented as SMILES (Simplified Molecular Input Line Entry System) strings. Because multiple SMILES strings can correspond to the same structure, and there is no universal approach for generating a canonical SMILES string, the strings were converted to InChI (International Chemical Identifier) keys [46] using RDKit to identify duplicate compounds.

From the unique set of small molecules, distinct unlabeled sets were derived for each of the three targets (hERG, Nav1.5, Cav1.2) using the following methodology. First, the ChEMBL unlabeled data were combined with the labeled data of each of the other two targets, collectively forming an extended unlabeled set. Subsequently, the extended unlabeled set underwent filtering to eliminate molecules overlapping with those in the target-labeled set of interest. Third, employing Tanimoto similarity, it was ensured that the extended set of unlabeled small molecules did not intersect, in terms of structural similarity thresholds, with the target external test sets (Eval-70 & Eval-60). This process yielded approximately 2 million unlabeled small molecules for each target.

A common issue encountered with large datasets is the presence of redundancy, persisting even after thorough data cleaning. Subsets of small molecules may cluster together due to shared properties or similar chemical structural components. Given training resource limitations, a strategic approach involves identifying a subset of data points that encapsulates the information inherent in the entire dataset. This subset can then be leveraged to train robust models that can learn further representations, thereby enhancing generalization. Submodular optimization has emerged as a promising solution for this NP-hard problem [47]. In this study, we employed submodular optimization using the 2048-bit extended connectivity fingerprints as a numerical representation for each compound. This method enabled the selection of an optimal representative subset comprising 100,000 (approximately 5%) unlabeled small molecules for each target. Regarding the submodular optimization algorithm used, we employed apricot [48] maximum coverage function using the default ‘two-stage’ optimizer, which integrates both a naïve greedy and a stochastic accelerated algorithm to rank molecules based on their gain with respect to the entire dataset. The remaining settings used the default configuration (refer to Supplementary Tables S1, S2, and S3 for the respective selected small molecules ranked by gain).

The full extended dataset of ∼2 million unique compounds is saved in an hdf5 small molecule library store and includes the following fields: InChI key, standardized SMILES string, compound source (refer to section 2.1 for all sources), ChEMBL identifier if the compound exists in this open access database, 1024-bit Morgan fingerprint, 2048-bit Morgan fingerprint, 881-bit PubChem fingerprints, 854 vector-length of preprocessed and standardized Mordred descriptors, and cardiotoxicity inhibition predictions for each of the three cardiac target using CtoxPred2 along with the model confidence scores. Consult Zenodo (https://zenodo.org/records/11066707) for access to the small molecule library store and a notebook demonstrating how to query it.

### 2.2. Molecular Features

The PyBioMed Python package [49] was used to compute molecular fingerprints. This process generated two types of fingerprints: extended connectivity fingerprints with a maximum diameter parameter of two (ECFP2) (a vector of 1024 ECFP fingerprint values) and PubChem fingerprints (a vector of 881 values).

The calculation of molecular descriptors was performed using the Mordred Python package [50]. We employed 2D descriptors only as these require fewer computational resources compared to 3D descriptors without sacrificing predictive performance [20][51][52][53], resulting in a total of 1613 descriptors. Preprocessing and feature selection were accomplished through a Scikit-Learn [54] pipeline consisting of four modules. First, a univariate imputer was employed to discard columns with no calculated values and replace missing values in other columns with the mean. Second, a standardization step was applied to remove the mean and scale the values to unit variance. Third, zero-variance features were removed. Finally, for any pair of highly correlated features (Pearson correlation above 0.95), one of them was randomly discarded, as such correlated features convey nearly identical information. Consequently, the unsupervised preprocessing procedure, which was performed on a combined labeled and unlabeled set of ∼2 million small molecules, reduced the feature space for all targets (i.e. hERG, Nav1.15, and Cav1.2) to 854 descriptors (refer to Supplementary Table S4 for the respective descriptor names). Afterwards, similar preprocessing steps were applied to the Eval-70 and Eval-60 test sets.

### 2.3. Model Building

Using a combination of 1905 fingerprints and 854 preprocessed descriptors as input features, we employed the SSL paradigm, particularly the pseudo-labeling technique, to build robust ML models. Pseudo-labeling leverages the concept of employing the model itself to assign artificial labels to unlabeled data [55][56]. More specifically, the approach involves the compilation of “hard” labels, which are determined by selecting the class with the highest probability output by the model. Next, only artificial labels with class probabilities exceeding a predetermined threshold are retained. The remaining data are discarded and re-evaluated in the next iteration to decide on a confident label assignment [57].

We utilized the self-training classifier available in Scikit-Learn. This estimator implementation is based on Yarowsky’s algorithm [58]. With this algorithm, a supervised model can act as a semi-supervised classifier, enabling learning from unlabeled data. To train the final model, the SSL meta estimator needs to be combined with another classifier, referred to as the base classifier. We opted for a RF as our base classifier. In each iteration, the base classifier predicts labels for the unlabeled samples and then adds a subset of the unlabeled data along with the artificial labels to the labeled dataset. Here, we included the 50 most confidently predicted samples per iteration for a maximum of 10 iterations, thus augmenting the training data with up to 500 previously unlabeled samples. The remaining hyperparameters used the default configuration. For hyperparameter tuning, we employed five-fold cross-validation (CV) with a predefined grid search space (Supplementary Table S7).

For each target (i.e., hERG, Nav1.5, Cav1.2), RF models were trained using two machine learning paradigms: supervised and semi-supervised learning. Under each paradigm, predictive performance was evaluated for three feature combinations: utilizing fingerprints only, descriptors only, or both fingerprints and descriptors combined. Hyperparameter tuning using five-fold CV within a predefined grid search space was used for all models.

Data pre-processing, cleaning, analysis, hyperparameter tuning, and model training were performed on a Spark cluster comprising one driver node (61GB RAM and 8 cores) and four worker nodes (each with 16GB RAM and 2 cores). Following model selection based on each feature combination, the top-performing models were evaluated on two external datasets. The final best models are denoted as CToxPred2-hERG, CToxPred2-Nav, and CToxPred2-Cav for the respective targets.

### 2.4. Statistical tests

We conducted two statistical analyses to assess model performance and compare SSL to supervised learning. First, we utilized a bootstrapping analysis on the prediction models using the test sets. Second, we employed McNemar’s test to compare the performance of the supervised and SSL classifiers. McNemar’s test [59], also known as “within-subject chi-squared” test, is a non-parametric statistical test for paired comparisons that can be applied to compare the performance of two machine learning classifiers. The test requires a categorical dependent variable with two categories (in our case blockers vs non-blockers), two models to compare, and performed on the same test set. The test itself is based on a version of the 2×2 confusion matrix [60], where each quadrant is referred to as A, B, C, and D. Due to the relatively small values of B and C in our tests, and the sum of both B and C being less than 50, we employed the recommended Binomial test to compute the exact p-value [60].

### 2.5. Evaluation Metrics

During the training process, the hyperparameters that yielded the best performance were then utilized to train the final model on the entire dataset. These models were subsequently evaluated on external test sets comprising 60% and 70% structurally dissimilar molecular compounds.

The performance of the models was assessed using multiple binary evaluation metrics, including accuracy (AC), sensitivity (SE), specificity (SP), F1-score (F1), correct classification rate (CCR), and Matthew’s correlation coefficient (MCC), as defined below:

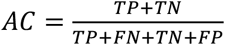

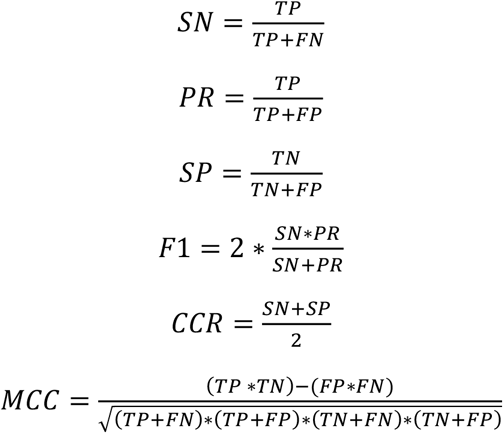

The final model selection was performed based on the F1 score.

### 2.6. Screened datasets

Using the trained CToxPred2 models, we performed an analysis of blockers identified from three distinct publicly available datasets: the complete labeled set encompassing approximately 25,373 unique molecules [62], all ChEMBL data extracted from the preprocessed extended library store [63], and a set comprising 1692 unique FDA-approved drugs sourced from the DrugCentral database [64].

The identification of blockers followed this process: if a molecule possessed an actual potency derived from assays as compiled in the labeled set, the label was assigned accordingly. For molecules not included in the labeled development set, CToxPred2 predictions were used instead, accompanied by the corresponding model probability confidence.

### 2.6. Graphical User Interface

To enhance the software usability for both technical and non-technical users, we incorporated the optimal models into a graphical user interface (GUI). Employing tkinter [61], the standard Python interface to the Tcl/Tk GUI toolkit (version 8.5 or higher), we developed a cross-platform GUI in Python. Tkinter comes bundled with most Python installations, including the standard library of Python itself, thus waiving the need for additional library installations. This GUI offers an interactive platform for analyzing small molecules under development, calculating eight key physicochemical properties using the quantitative estimation of drug-likeness (QED) [26][27] module in RDKit, alongside 2D structure rendering, cardiotoxicity predictions, and confidence prediction measures, serving as an indicator of model uncertainty. Notably, the application operates on users’ premises, ensuring the privacy of their drug data without the need for online exposure to public websites or third-party software (Supplementary Figure S1).

## 3. Results

### 3.1. A Comprehensive SSL Dataset of Cardiac Ion Channel Blockers

The dataset presented here serves as a valuable resource for researchers operating within the domain of drug discovery to conduct in-depth analyses and further studies in the field. The database comprises three components: an extensive large curated small molecule store, a large labeled cardiotoxicity dataset, and a representative unlabeled dataset sampled using submodular optimization utilized for SSL.

The extensive small molecule store comprises approximately 2 million compounds, presenting a significant challenge for leveraging such a massive dataset for cardiotoxicity predictions. First, there is a significant disparity and imbalance between labeled and unlabeled compounds, with only a few hundred or thousand labeled molecules compared to unlabeled ones. Second, this database surpasses previous training datasets in scale by orders of magnitude, leading to potential issues with excessive training times and data volume challenges. Some of these challenges include potentially redundant information which may not contribute significant signal to the ML models. Consequently, we have employed submodular optimization as a principled approach to efficiently address redundancy while preserving data integrity. By strategically selecting a subset of unlabeled compounds, this ensures that the dataset remains representative and diverse. The selected data represents a 5% sampled subset from the cleaned and unique set of all collected 2 million unlabeled small molecules. Employing Tanimoto similarity, it was ensured that the 100,000 unlabeled small molecules sampled for each target ion channel did not overlap, in terms of structural similarity thresholds, with the two external test sets (Eval-70 & Eval-60; Supplementary Figure S2). Importantly, this sampling retained the independence and identically distributed (iid) assumption between the training and test sets, which is crucial for ML model development (Figure 1).

**Figure 1.**
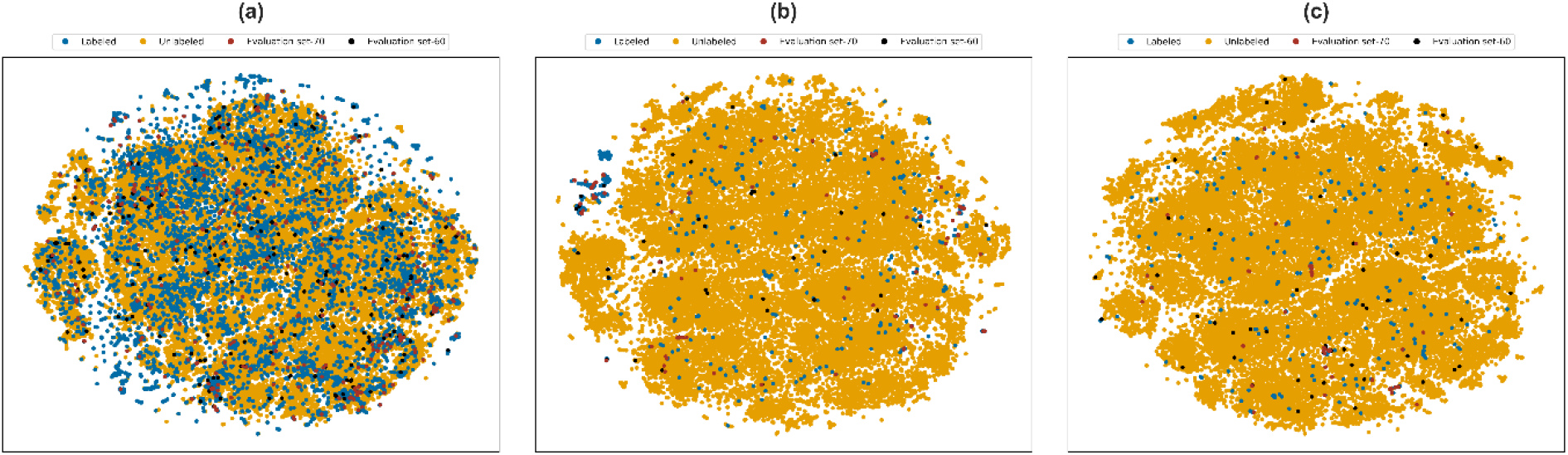
t-SNE visualizations showing the distributions of the labeled and unlabeled molecules in the development set and the two external test sets (Eval-60 and Eval-70) for (a) hERG (b) Nav1.5 and (c) Cav1.2.

As a special remark for Nav1.5, we observed that a significant number of labeled compounds in the structural molecular space was clustered together away from the unlabeled data, which contributed to some bias in the Nav1.5 labeled set. Consequently, when only training on labeled data, the model will overfit to these specific molecular structures, limiting its ability to generalize to the much more diverse chemical space. On the other hand, SSL will be able to alleviate that bias and provide more generalizing models.

### 3.2. Evaluation of SSL For Machine Learning of Cardiotoxicity

Using both fingerprints and descriptors combined as input features to our predictive models, we examined the performance gain of SSL compared to supervised learning across all three targets (Table 1). In general, SSL improves performance on the most challenging task (Eval-60). This benefit of SSL holds for different input features as well, such as only using fingerprints or descriptors (Supplementary Tables S12, S13, & S14). If we pick the case of Nav1.5 and Cav1.2, SSL exhibited performance gains across all test sets, but more noticeably for structurally dissimilar evaluation sets, highlighting the positive influence of leveraging unlabeled data, especially in training sets with limited labeled data. Regarding hERG specifically, the impact of unlabeled data used during training showcased a significant enhancement in model performance, but particularly evident in evaluation sets exhibiting large dissimilarities when employing SSL, while it remained relatively unchanged for hERG-70. This outcome can be attributed to the size of the labeled training set for this target and the distribution of these compounds within the molecular structural space, which was relatively extensive and covering a wider structural space (Figure 1a).

**Table 1.** Cardiotoxicity prediction performance on the three ion channels combining fingerprints and descriptors as input features to a random forest model following either a supervised or a semi-supervised learning approach. The evaluation was conducted on two external test sets, Eval-70 and Eval-60, showing an increased generalization of the models on structurally dissimilar test sets when using SSL.

| ML Paradigm | Model Input Features | Eval-70 |  |  |  |  |  | Eval-60 |  |  |  |  |  |
| --- | --- | --- | --- | --- | --- | --- | --- | --- | --- | --- | --- | --- | --- |
|  |  | AC | F1 | SN | SP | CCR | MCC | AC | F1 | SN | SP | CCR | MCC |
| Supervised learning | RF-Cav | 88.9 | 91.6 | 94.2 | 79.3 | 86.8 | 75.5 | 67.7 | 74.4 | 70.7 | 61.9 | 66.3 | 31.5 |
|  | RF-Nav | 84.5 | 88.9 | 90.7 | 71.1 | 80.9 | 63.5 | 65.6 | 77.1 | 77.1 | 31.2 | 54.2 | 8.3 |
|  | RF-hERG | 82.0 | 84.9 | 90.5 | 71.3 | 80.9 | 63.7 | 70.0 | 72.9 | 80.8 | 59.2 | 70.0 | 41.0 |
| Semi-supervised learning | CtoxPred2-Cav | 91.4 | 93.6 | 98.1 | 79.3 | 88.7 | 81.3 | 74.2 | 78.9 | 73.2 | 76.2 | 74.7 | 47.1 |
|  | CtoxPred2-Nav | 85.9 | 90.2 | 94.8 | 66.7 | 80.8 | 66.4 | 68.8 | 79.6 | 81.2 | 31.2 | 56.2 | 13.1 |
|  | CtoxPred2-hERG | 82.0 | 85.1 | 91.7 | 70.0 | 80.8 | 63.8 | 72.8 | 75.2 | 82.4 | 63.2 | 72.8 | 46.5 |

To further quantify the benefits of SSL, we conducted two statistical analyses. First, a bootstrapping analysis was performed on the prediction models using the test sets (Supplementary Table S5). Second, we employed McNemar’s test to compare the performance of the supervised and SSL classifiers. Supplementary Table S6 displays the 2×2 McNemar matrix comparing the two models, from Table 1, for each target. While none of these tests achieved p-values below a significance threshold of 0.05, they do show lower p-values when moving from the Eval-70 to the Eval-60 test sets, suggesting that the classifiers are not equivalent and SSL models are indeed more performant and generalize better to highly different compounds than their purely supervised counterparts.

Additionally, we conducted a comparative analysis of our final method (CToxPred2) with the state-of-the-art neural network-based CardioGenAI tool [65], both trained and evaluated on the same datasets (Supplementary Table S8). This evaluation indicated CardioGenAI’s competitive performance on the Eval-70 test sets compared to CToxPred2 (Figure 2a). However, when evaluating the performance of both methods on the strictly dissimilar test sets (hERG-60, Nav-60, and Cav-60), CToxPred2 increasingly outperformed CardioGenAI, indicating its increased generalization performance, likely due to its semi-supervised training (Figure 2b). This is especially notable for the Nav-60 dataset, for which the labeled data exhibits a bias towards a specific family of molecules, as previously described. This suggests potential overfitting and a lack of generalization capability for CardioGenAI compared to CToxPred2.

**Figure 2.**
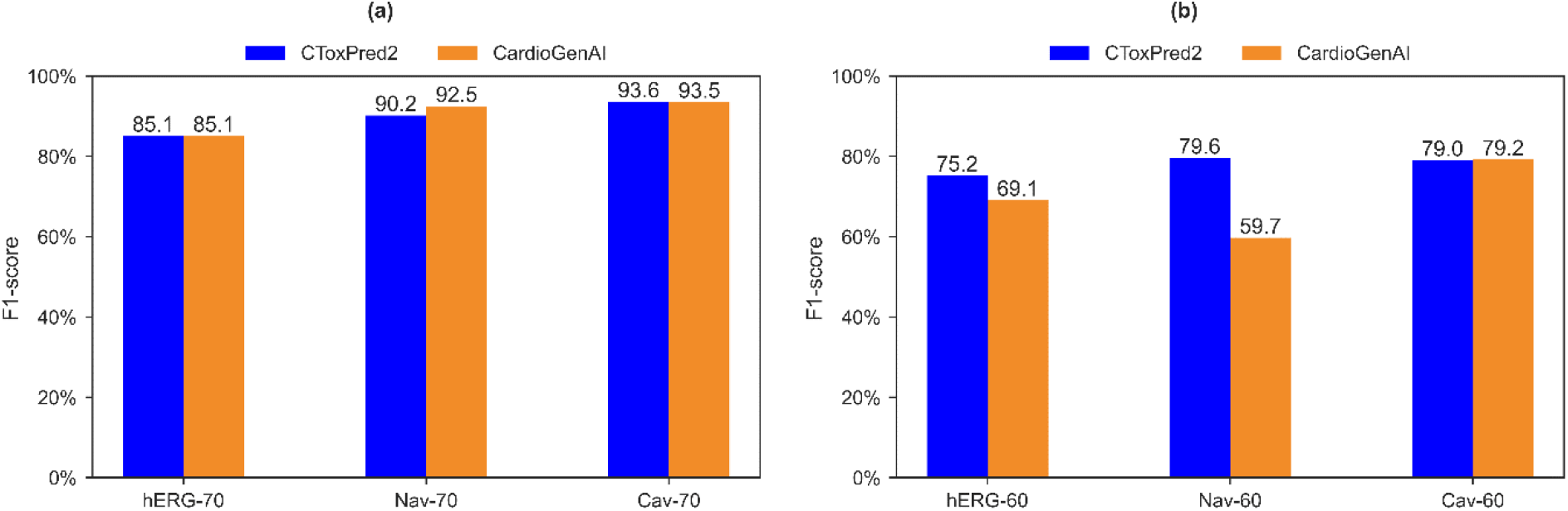
Evaluation of the CToxPred2 performance compared to CardioGenAI [65], trained and tested using the same data, where (a) is performance on Test-70 and (b) performance on Test-60.

### 3.3. From Prediction to Discovery: Unveiling Confidently Identified Novel Molecular Blockers

Using the CToxPred2 models, we conducted an analysis of blockers identified from three distinct publicly available datasets. The first dataset under examination is the complete labeled set, encompassing 25,373 unique molecules. Notably, 54.5% of these molecules exhibit hERG blocking activity, while only 9.3% and 2.6% demonstrate Nav1.5 and Cav1.2 blocking activity, respectively (Supplementary Figure S3). This highlights the predominant emphasis placed on hERG in past assessments of drug cardiovascular toxicity. Within this set, 36 molecules are classified as potent blockers capable of inhibiting all three cardiac ion channels (Figure 3a).

**Figure 3:**
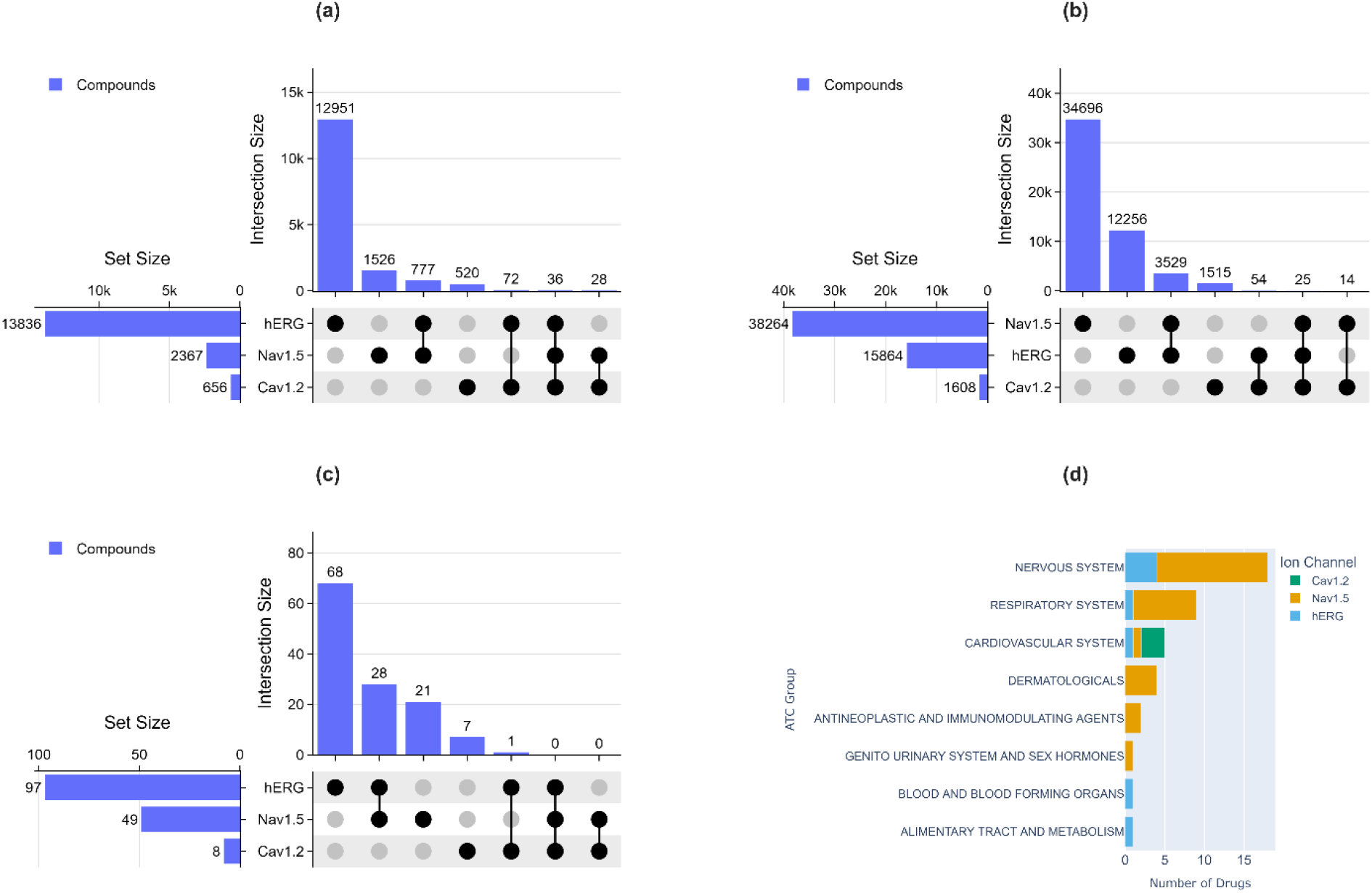
Analysis of blockers within three datasets. (a) Labeled set derived from potencies reported in assays, or predicted if no assay available. (b) ChEMBL, for both potency-based and confidently predicted blockers. (c) FDA-approved confidently predicted blockers. (d) Distribution of identified FDA-approved drugs (not included in training process) exhibiting inhibition activities of the three ion channels based on their ATC group.

Next, we used CToxPred2 to discover potential cardiac ion channel blockers in two external, unlabeled datasets: ChEMBL and FDA-approved drugs.

Employing a confidence threshold of 90% or higher during the screening process unveiled a significant number of Nav1.5 blockers, constituting 2.0% of the ChEMBL dataset. In contrast, fewer blockers were predicted for hERG and Cav1.2, with respectively 0.8% and 0.1% of the dataset (Figure S4). Overall, 25 compounds were identified as potent inhibitors capable of blocking all three targets within ChEMBL (Figure 3b).

Additionally, we evaluated the cardiac ion channel inhibition of drugs used in the market by screening 1692 unique FDA-approved drugs sourced from the DrugCentral database [64]. This resulted in 97 drugs predicted as hERG blockers, 49 as Nav1.5 blockers, and 8 as Cav1.2 blockers (Figure 3c). Subsequent analysis revealed an overlap between the FDA-approved drugs analyzed and the training set. After excluding compounds present in the development data, the models still indicate “in silico” that there is a potential for cardiotoxicity of the following compounds: 8 actively used drugs were still predicted as potential hERG blockers, 29 as potential Nav1.5 blockers, and 3 as potential Cav1.2 blockers. While an interesting observation, to confirm these inhibition side effects, experimental observations are necessary, as numerous factors can influence the in vivo outcomes.

To gain further insights into the operation and prevalence of these drugs, we queried them against DrugBank [66][67] to obtain their names, FDA approval status, and ATC codes corresponding to their target systems (Supplementary Tables S9, S10, and S11). Upon analyzing the target systems for which these drugs are designed, we observed that drugs targeting the nervous system are more likely to exhibit cardiovascular inhibition properties (Figure 3d). Therefore, we recommend thorough testing of the inhibition capabilities of such drugs and evaluating their modes of administration and uptake to avoid deleterious cardiac side effects.

## 4. Conclusion

Drug-induced cardiotoxicity, a leading cause of drug withdrawals, poses significant challenges in pharmaceutical research. Early prediction of drugs’ pharmacokinetic properties is crucial to mitigate such risks. While conventional ML has shown high promises, the scarcity of ground truth, annotated toxicity data hampers model performance. Unlabeled data, often overlooked, holds immense potential to unveil hidden patterns within chemical and biological systems. By tapping into this resource, researchers can refine their prediction models to better generalize to unseen data. In this study, we mined a large unlabeled dataset of small molecules and applied SSL to enhance the performance of cardiotoxicity liability prediction. Our results demonstrate that SSL improves the robustness and generalizability of these models. However, it is important to acknowledge that the observed improvements in performance are not that large and lack strong statistical significance. This highlights the need for future research to understand which compounds are the most relevant to compile larger unlabeled training datasets. Currently, we have addressed this by using submodular optimization to maximize the molecular structural diversity of the training data, while ensuring the dataset size remains manageable. Next, we have developed a cross-platform, user-friendly GUI to enable easy access to our models, allowing users to conveniently assess the blocking property of their own compounds. Finally, we applied our trained models to assess the cardiac ion channel inhibition behavior of drugs currently active on the market, revealing that several drugs may exhibit potential blocking activity. As discussed earlier, this remains an in silico study that requires further investigation. To confirm such inhibitory side effects, experimental observations are essential, as many other factors influence in vivo outcomes. It is also worth noting that in silico predictions should be used as supporting tools, providing an additional layer of information for drug development pipelines. They do not have the authority to undermine an approval process that typically involves clinical trials.

## Data and Software Availability

CToxPred2 is available as an open-source Python GUI-based tool and can be called from a notebook. It uses Pybel (version 0.13.2) [68], Open Babel (version 3.1.1) [69], and PyBioMed (version 1.0) [49] to compute PubChem & ECFP2 fingerprints; Mordred (version 1.2.0) [50] to calculate molecular descriptors; RDKit (version 2023.9.5) [70] for chemical structure information retrieval, used for example in the graph representation construction and other tasks; Scikit-Learn (version 1.3.1) [54] for pipeline data preprocessing, evaluation metric calculations, and SSL trained models; PyTorch (version 1.13.1) [71] libraries to maintain the supervised deep learning models previously deployed in CToxPred; and NumPy (version 1.23.5) [72], SciPy (version 1.11.4) [73], and Pandas (version 2.0.3) [74] for scientific computing. Matplotlib (version 3.6.2) [75], Seaborn (version 0.13.2) [76], and Plotly (version 5.21.0) [77] were used for data visualization. Data analysis was performed using Jupyter notebooks [78].

CToxPred2 is available as open source under the permissive MIT license on GitHub at https://github.com/issararab/CToxPred2. Analysis notebooks to reproduce the presented results are also available in the same repository at https://github.com/issararab/CToxPred2/blob/main/notebooks/analysis_notebook.ipynb. All data used in this study are freely available through Zenodo for long-term archival at https://zenodo.org/records/8359714 & https://zenodo.org/records/11066707. The data are also available on the CToxPred2 GitHub repository, alongside precomputed results of the screened datasets: https://github.com/issararab/CToxPred2/tree/main/data.

## Notes

https://zenodo.org/records/11066707

https://zenodo.org/records/8359714

https://github.com/issararab/CToxPred2

### Competing Interest Statement

The authors have declared no competing interest.

