## Supplementary Figures for "Semi-Supervised Learning to Boost Cardiotoxicity Prediction by Mining a Large Unlabeled Small Molecule Dataset"

### Supplementary Figures to “Semi-supervised Learning to Boost Cardiotoxicity Prediction by Mining a Large Unlabeled Small Molecules Dataset”


Issar Arab, Kris Laukens , Wout Bittremieux

Department of Computer Science, University of Antwerp, 2020 Antwerp, Belgium

Biomedical Informatics Network Antwerpen (biomina), 2020 Antwerp, Belgium


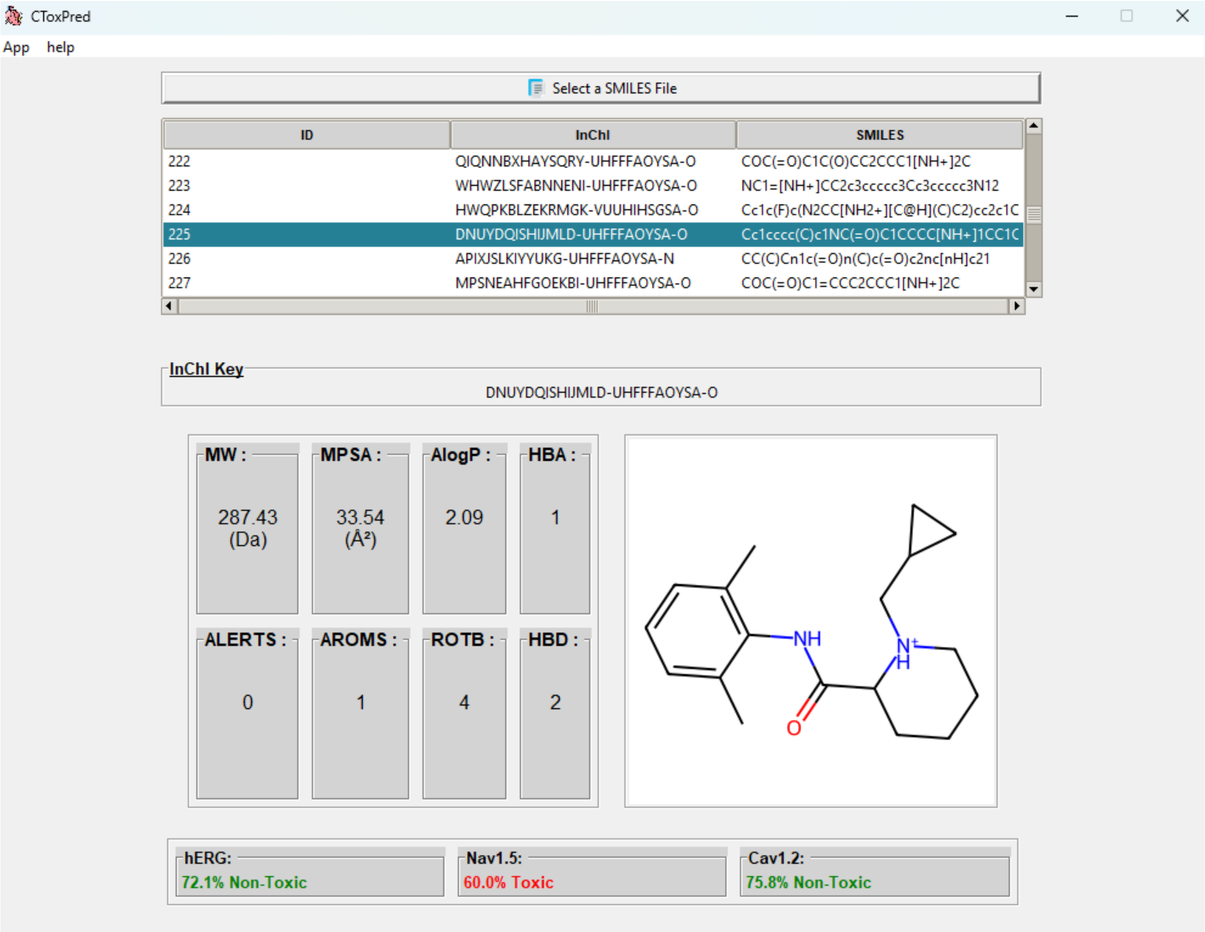


**Supplementary Figure S1:** Home page of CToxPred2, a cardiotoxicity prediction software. Using the select button, users can navigate their devices and choose a file containing SMILES strings. The file extension must be either ".smi" or ".smiles". Once all metrics are computed, users can interactively select the molecule of interest from the list. They can then view the computed eight physicochemical properties, the 2D structure, and the predictions for cardiotoxicity, along with confidence scores provided by the models.

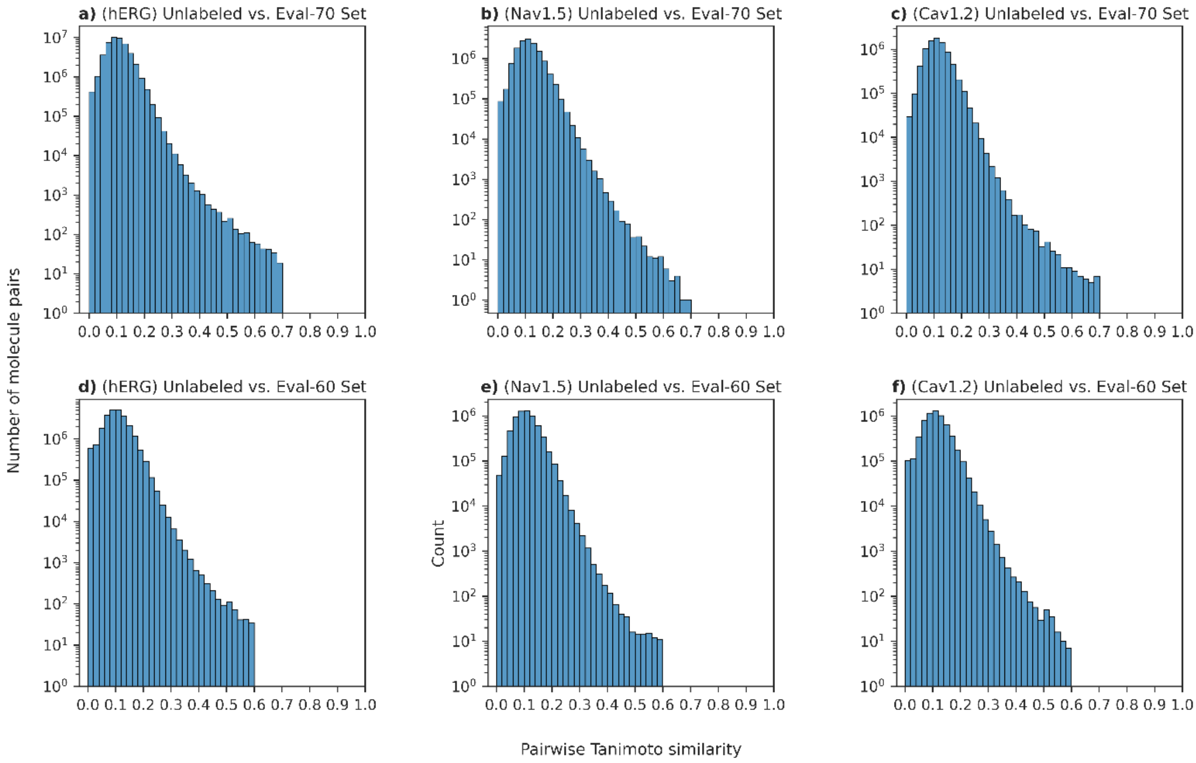


**Supplementary Figure S2:** Distribution of the pairwise Tanimoto similarity for each molecule in the **(a)** hERG unlabeled set with the ones in the evaluation set hERG-70, **(b)** Nav1.5 unlabeled set with the ones in the evaluation set Nav-70, **(c)** Cav1.2 unlabeled set with the ones in the evaluation set Cav-70, **(d)** hERG unlabeled set with the ones in the evaluation set hERG-60, **(e)** Nav1.5 unlabeled set with the ones in the evaluation set Nav-60, **(f)** Cav1.2 unlabeled set with the ones in the evaluation set Cav-60.

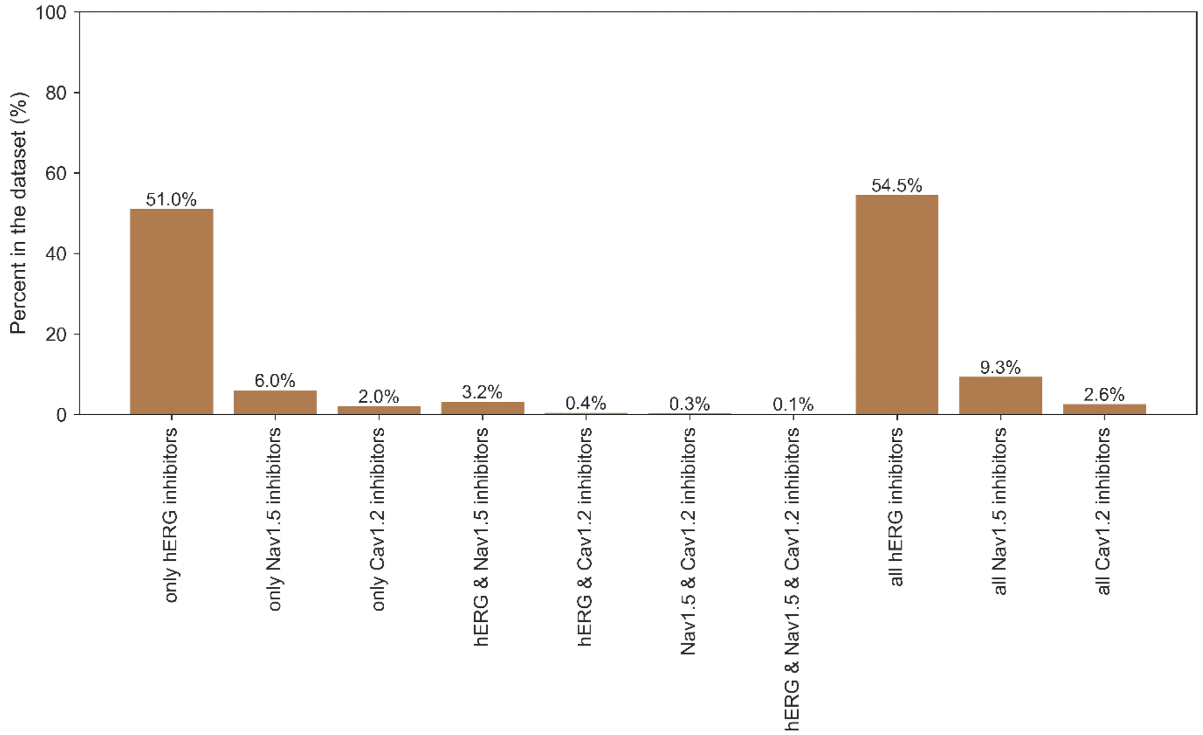


**Supplementary Figure S3:** The distribution of blockers within the labelled small molecules database, expressed as a percentage. These blockers are identified from the potency values obtained through conducted assays.

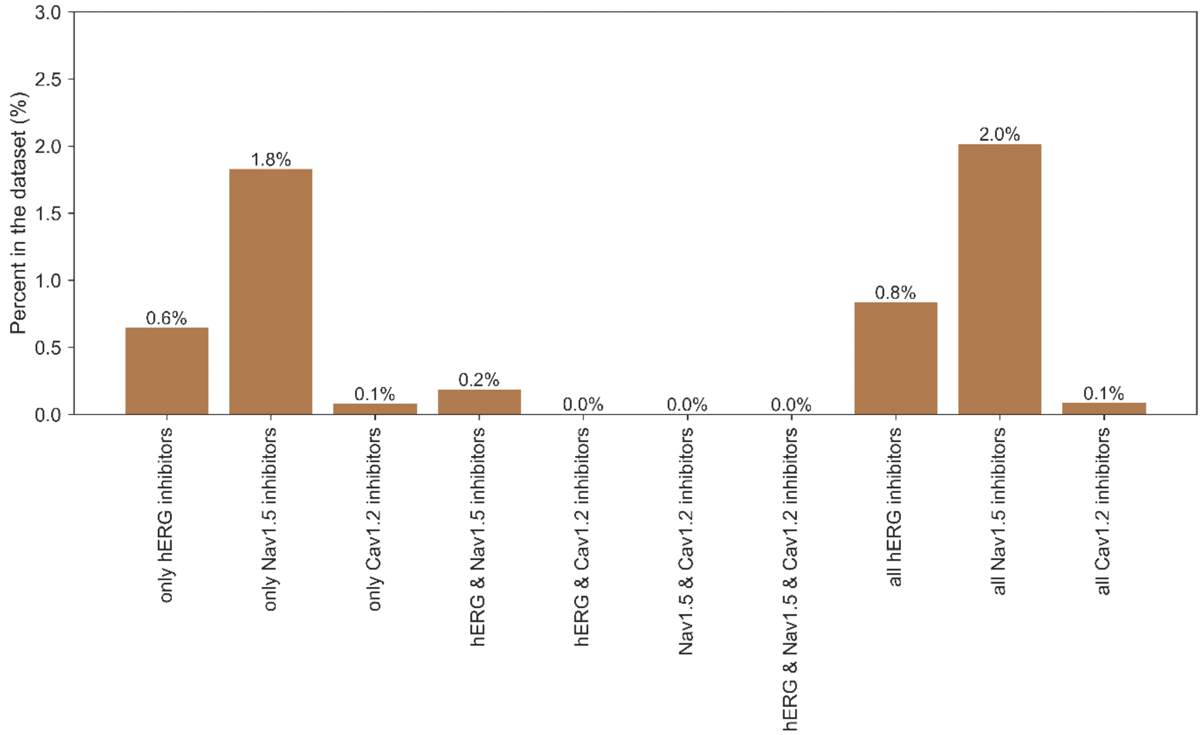


**Supplementary Figure S4:** The distribution of confident existing blockers within the ChEMBL small molecules database, expressed as a percentage. These blockers are either exact blockers, identified directly from potency values obtained through conducted assays, or predicted by the CToxPred2 model with a confidence score of 90% or higher.
